# Multiplexed computations in retinal ganglion cells of a single type

**DOI:** 10.1101/080135

**Authors:** Stephane Deny, Ulisse Ferrari, Emilie Mace, Pierre Yger, Romain Caplette, Serge Picaud, Gašper Tkačik, Olivier Marre

## Abstract

In the early visual system, cells of the same type perform the same computation in di↵erent places of the visual field. How these cells code together a complex visual scene is unclear. A common assumption is that cells of the same type will extract a single stimulus feature to form a feature map, but this has rarely been observed directly. Using large-scale recordings in the rat retina, we show that a homogeneous population of fast OFF ganglion cells simultaneously encodes two radically different features of a visual scene. Cells close to a moving object code linearly for its position, while distant cells remain largely invariant to the object’s position and, instead, respond non-linearly to changes in the object’s speed. Cells switch between these two computations depending on the stimulus. We developed a quantitative model that accounts for this effect and identified a likely disinhibitory circuit that mediates it. Ganglion cells of a single type thus do not code for one, but two features simultaneously. This richer, flexible neural map might also be present in other sensory systems.

## Introduction

A major challenge of the visual system is to extract meaningful representations from complex visual scenes. Feature maps, where the same computation is applied repeatedly across different sub-regions of the entire visual scene, are essential building blocks for this task, for both sensory networks (Fitzpatrick and Ulanovsky, 2014; Ohki et al, 2005) and artificial vision systems (LeCun et al, 2015). Ganglion cells, which form the retinal output, can be divided into different types (Wassle and Boycott, 1992; Devries et al, 1997; Field et al, 2010; Baden et al, 2016). In the classical view of retinal function, cells of the same type extract a single feature from the visual scene and generate a feature map that is then sent to the brain (Azeredo da Silveira R and Roska, 2011). This “one type = one feature” view is well illustrated in the retina when objects move across the visual field at constant speed. In this case, previous work has shown that a single type indeed represents a single feature of the scene (Berry et al, 1999; Vaney et al, 2012; Leonardo and Meister, 2013; Trenholm et al, 2013).

However, processing by ganglion cells also depends on the visual context (Shapley and Enroth-Cugell, 1984; Smirnakis et al, 1997; Geffen et al, 2007; Farrow et al, 2013; Tikidji-Hamburyan et al, 2015), so that feature extraction will be influenced by the global parameters of the visual scene, e.g., by its luminance and contrast. Furthermore, ganglion cell activity can be modulated by stimulation outside of the cells’ classically-defined receptive fields (McIlwain, 1964; Roska and Werblin, 2003; Passaglia et al, 2001, 2009; Marre et al, 2015), implying that feature extraction may not be entirely local, especially when presented with complex, dynamical stimuli. As a result, it is not clear how irregular trajectories of moving objects, which are ubiquitous in natural scenes (Eizenman et al, 1985; Branson et al, 2009), are represented by ganglion cells of the same type.

Here we show that a single ganglion cell type extracts simultaneously two very different features from a visual scene composed of irregularly moving bars. Within a homogeneous population of fast OFF ganglion cells recorded simultaneously, cells whose receptive field center overlapped with an object performed a linear computation that was highly sensitive to the position of the object. In contrast, cells of the same cell type that were far from any moving object responded nonlinearly to fast motion, and were largely invariant to the exact position of distant objects. Individual cells switched from one computation to the other when their receptive field center was stimulated. We constructed a model that quantitatively accounted for these findings, and determined that the observed scheme of distal activation is implemented by a disinhibition circuit of amacrine cells.

## Results

### Cells of a single cell type respond to very distant moving objects

We recorded large ensembles of ganglion cells from the rat retina using a micro-electrode array of 252 electrodes (Marre et al, 2012; Yger et al, 2016). We measured the receptive field center of each cell with binary checkerboard noise. To separate ganglion cells into different types, we displayed several stimuli (full field flicker, drifting textures) and grouped together cells with similar responses (see Methods). In the following, we focus on a single group composed of well-isolated fast OFF cells. Their responses to spatially uniform stimuli were nearly identical (fig. 1A), and their receptive fields clearly tiled the visual space (fig. 1B).

**Figure 1:**
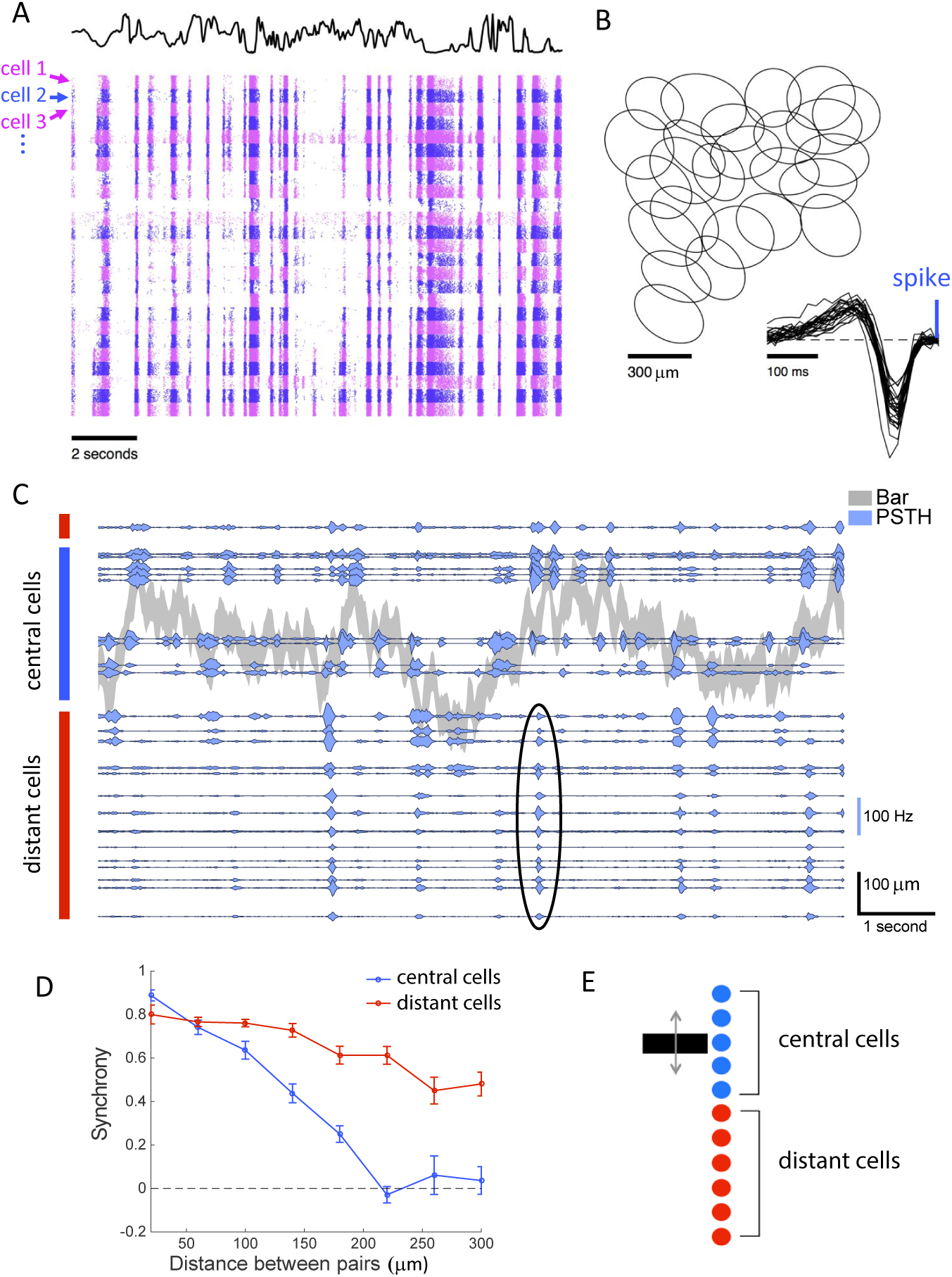
A single cell type responds synchronously to distant moving objects. **A:** Raster of 25 cells of the same type responding to a full field uniform flicker. Each line corresponds to a repeat of the stimulus, and each cell is indicated by a different color (alternating pink and blue). The black curve indicates the light intensity of the flicker over time. **B:** Receptive fields of a population of ganglion cells of the same type. Each ellipse represents the position and shape of the spatial receptive field associated with one cell (1-SD contour of the 2D Gaussian fit to the spatial profile of the RF). Inset: temporal profiles of the receptive fields of the same cells. **C:** PSTHs of multiple ganglion cells responding to repeated presentations of a randomly moving bar. Gray shade: position of the bar as a function of time (shade width corresponds to the bar width). Blue traces: PSTHs of individual ganglion cells, with baselines positioned to scale relative to the bar. Blue and red vertical rectangles indicate central and distant cells, respectively. Black ellipse shows an example synchronous firing event of the distant cells. **D:** Average SE cross-correlation between PSTHs of pairs of cells, as a function of their pairwise distance measured along the bar motion axis. Curves shown separately for cells whose receptive field center either was (blue) or was not (red) stimulated by the bar. **E:** Schematic diagram shows central cells (blue) and distant cells (red) that respond synchronously.

We then displayed a bar moving randomly over the visual field. This dark bar over a gray background was animated by a Brownian motion with a feedback force to keep the bar positioned over the array. As expected, ganglion cells whose receptive field center overlapped with the bar position responded reliably to a repeated trajectory, as shown by their PSTHs in fig. 1C. More surprisingly, reliable responses were also elicited in cells whose receptive field centers were far away from the bar. The receptive field center diameter was on average 287 ± 23 *µ*m (mean ± SD, n=25), and cells as far as 670*µ*m from the closest bar position responded to the moving bar.

These distant cells fired synchronously to the moving bar, largely independently of the location of their receptive field, while central cells did not. Central cells were only synchronous when they were very close to each other. The mean cross-correlation between the responses of pairs of central cells was 0.02 ± 0.04 (mean ± SEM, n = 20 pairs) for cells separated by more than 200 *µ*m along the axis perpendicular to the bar. In comparison, distant cells remained synchronous over large distances (fig. 1D). The mean cross correlation was 0.53 ± 0.03 (mean ± SEM, n = 35 pairs, Pearson correlation r) for distant cells separated by more than 200 *µ*m. This distant activation had a profound effect on the structure of the retinal activity: while the bar covered a region 0.4 mm wide, ganglion cells were activated over an area wider than 1.4 mm (fig. 1E).

### Linear computation inside and non-linear computation outside of the receptive field center

We asked if the observed ganglion cell responses to motion outside their receptive field centers could be explained by standard models of the retina. We fitted a Linear-Non-linear-Poisson (LN) model (fig. 2A) to the response of each cell to non-repeated trajectories of the moving bar.

**Figure 2:**
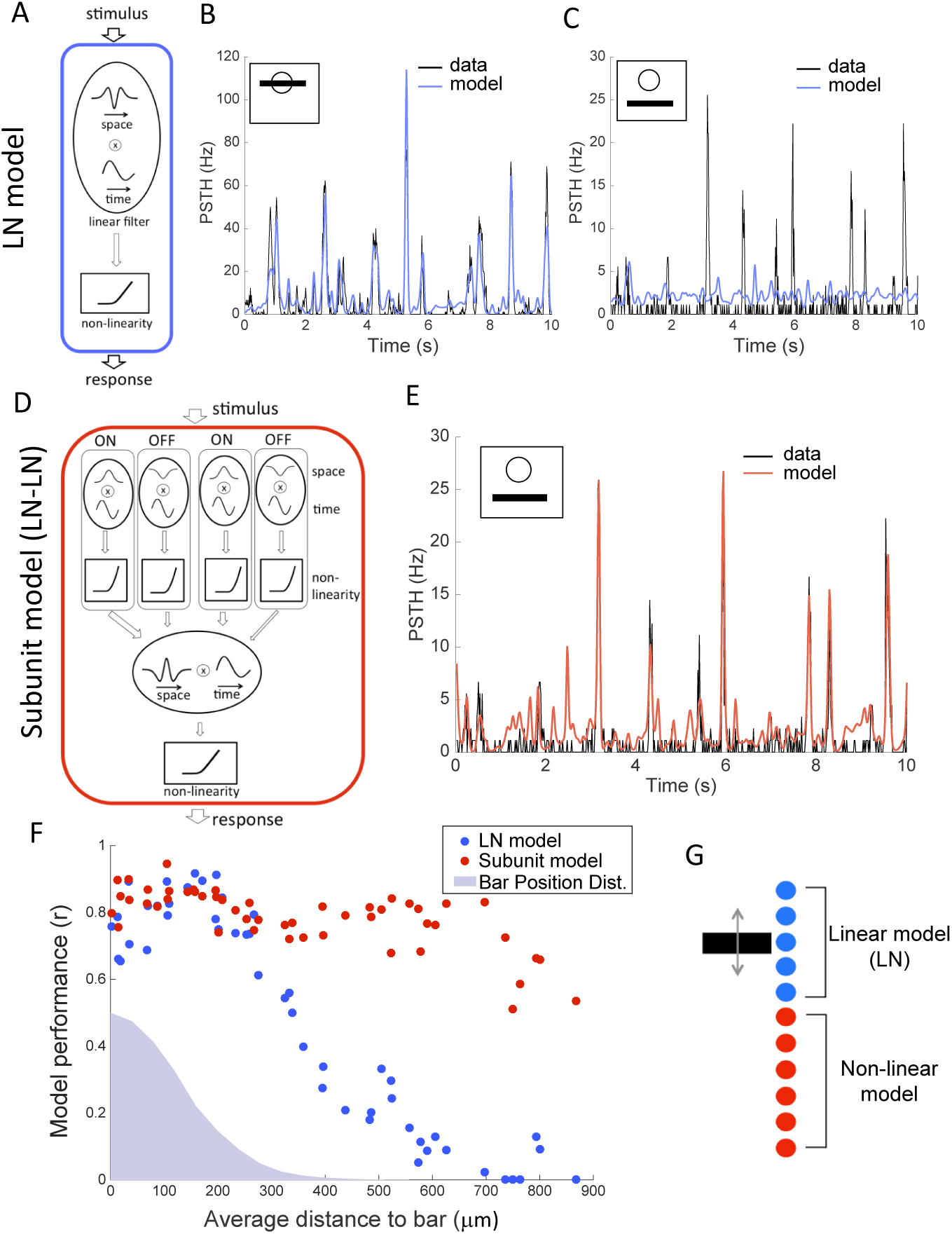
OFF ganglion cells perform a linear computation in their receptive field center, and a non-linear computation in the surround. **A:** Schematic of the LN model, composed of a linear filter and static non-linearity. **B:** Response (PSTH, gray) of a ganglion cell whose receptive field center is stimulated by the bar, is predicted by the LN model (blue). r = 0.89. **C:** Response (PSTH, gray) of the same ganglion cell when the bar is far from the receptive field center, is not predicted well by the LN model (blue). r = 0.02. **D:** Schematic of the subunit model, composed of a first stage (each subunit linearly filters the stimulus and applies a static nonlinearity), followed by weighted linear pooling and a second non-linearity. **E:** Response (PSTH, gray) of the same ganglion cell (as in A and B) to distant stimulation is predicted well by the subunit model (red). r = 0.83 **F:** Performance of the LN (blue) and subunit (red) models in predicting ganglion cell responses, as a function of the distance of the cell to the bar. Blue shade: position distribution of the bar. **G:** Schematic showing that cells whose receptive field center is on top of the moving bar perform a linear computation while distant cells perform a non-linear computation.

To test it, we repeated the same bar trajectory 54 times and compared predictions of the model with the measured PSTH for each cell. When the bar was moving close to or inside the receptive field center of the cell, the LN model predicted very well the response to the repeated sequence (r =0.79 ± 0.02, n = 25, fig. 2B). However, for cells that were distant from the bar, the same LN model failed at predicting their responses (fig. 2C, r =0.12 ± 0.02, n = 19). Performance was much lower (p 10^−25^, two-sample t-test), and could not be explained by a decrease in the reliability of the response (the ratio of explainable variability predicted by the model was 13% ± 2%, n = 19, see methods). This low performance was obtained despite the fact that we fitted the LN model directly on the responses to the distant bar. The low performance was also not due to any intrinsic property of these cells, but was related to the distance between the bar and the receptive field. When we displayed the moving bar in different locations, the same cells that were previously not predicted by the LN model (r =0.12 ± 0.02, n = 19), with RFs far from the bar, were predicted very well by a LN model when the bar was displayed inside their receptive field center (r =0.79 ± 0.02, n = 19; p ≤ 10^−13^, paired-sample t-test). In summary, the LN model was a good model for stimuli inside the receptive field center, but not outside.

To improve the prediction, we considered a model with two stages of processing that implements a non-linear summation within its receptive field (Victor, 1988; Gollisch and Meister, 2008; McFarland et al, 2013; Freeman et al, 2015; Vintch et al, 2015). The first stage was composed of many stereotyped subunits that convolved the stimulus with a linear filter and rectified the output to eliminate negative values. There were two types of subunits, ON and OFF, designed to mimic bipolar cell processing. The subunits of the same type were identical except that the linear filters were centered at different locations, such that they tiled the visual field. In the second stage, the outputs of the subunits was pooled together linearly in a weighted sum and then rectified to predict the firing rate (fig. 2D). To fit the model to data, we kept the first stage fixed and fitted the subunit weights in the second stage of the model (see methods for details).

This model predicted very well the responses of distant ganglion cells to a repeated random trajectory (r =0.73 ± 0.02, n = 19 fig. 2E). Performance was high for all distances of the receptive field to the bar (fig. 2F), demonstrating that the subunit model robustly captured responses that were not predicted by the LN model. Since the subunit model is a generalization of the LN model, it performed also well for center stimulation: in this case, rectified subunits were summed in the second stage of the model such that the net effect of a stimulus in the center was linearized (Werblin, 2010).

Our results showed that a population of cells of the same type extracted simultaneously two features from a single moving object. Cells whose receptive field centers overlapped with the object performed a linear computation on the stimulus, well recapitulated by an LN model. Distant cells performed a non-linear computation that was captured by a more complex subunit model described above. Therefore responses to a distant moving bar could not be simply explained by using a broader linear filter within the LN model framework. Taken together, these findings show that two radically different computations, performed on the same stimulus, can coexist within a population of ganglion cells of a single type (fig 2G).

### Switching between two modes of computation

Since the cells perform distinct computations in their center and in their distant surround, we studied how these computations interact when both center and surround are stimulated at the same time. What happens to distant cells if another bar is simultaneously shown inside their receptive field center? One possibility is that center and distant responses are simply added, so that the response to two moving bars would be the sum of the responses to each bar presented separately.

To test this, we displayed two bars moving randomly, with distinct trajectories, in two different locations. The distance between the bars’ average positions was 600 *µ*m. We also displayed each bar in isolation, at the same location and animated by the same trajectory as in the combined bar stimulus. We found that the response to the two bars was not equal to the sum of the individual responses to each bar presented separately (fig. 3A). Instead, if a bar was moving inside the receptive field center of the cell, the response to the distant bar was suppressed, while the distant bar exerted a negligible effect on the response to the central bar. Specifically, when one of the bars was moving inside the receptive field of a cell, the response to the combined bar stimulus was highly similar to the response to the single central bar (r =0.91 ± 0.01, n = 13). The residual response to the distant bar in the presence of simultaneously presented central motion correlated poorly with the response to the distant bar alone, and this discrepancy could not be explained by noise (fig. 3B, see methods for details).

**Figure 3:**
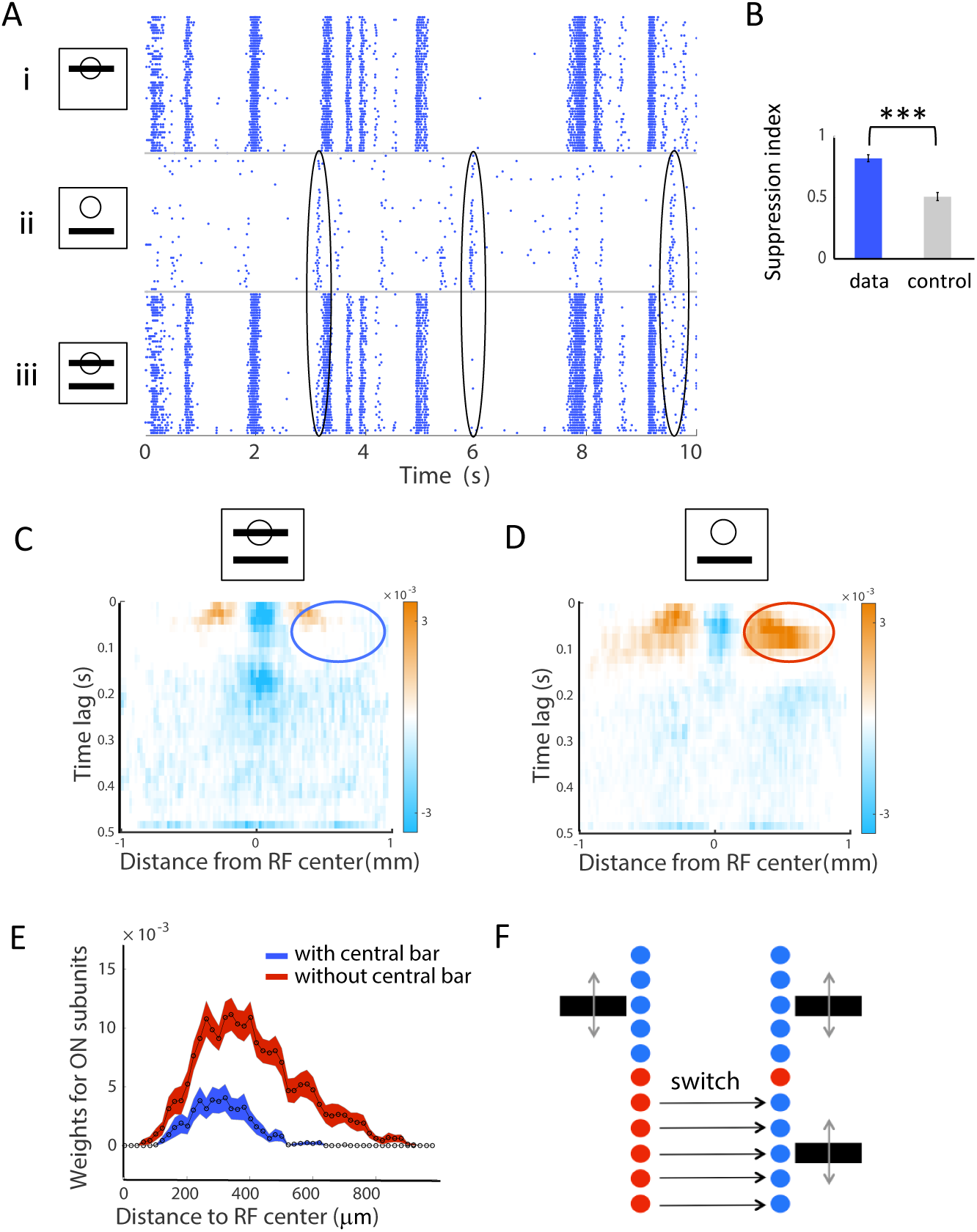
Suppression of distant responses by a bar moving inside the receptive field center. **A:** Response raster of a single cell to a moving bar presented in and/or outside of its receptive field center. Each dot is a single spike from the recorded cell. Each line corresponds to a different repetition of the same stimulus. i: bar moving inside the receptive field center. ii: bar moving outside the receptive field center. iii: both bars displayed together. Black ellipses indicate examples where the response to the distant bar is strongly suppressed by the stimulus inside the receptive field center. **B:** Suppression index for real cells (data, blue) and for the additive model (control, gray); see text and methods for details. **C, D:** Spatio-temporal distribution of the ON-subunit weights for the second stage of the subunit model, averaged over all cells. **C:** combined bar stimulus. **D:** a single distant bar stimulus. Blue and red ellipses show the reduction of the weights in the surround in the presence of central bar motion. **E:** ON-subunit weights summed over all times lags and averaged over all cells, as a function of distance to the receptive field center. Blue: combined bar stimulus, red: a single distant bar stimulus. **F:** Schematic showing how cells switch the mode of computation when a bar is displayed within their receptive field center.

To quantify further the observed suppression, we fitted the subunit model for the three stimuli separately (bar 1, bar 2, two bars). We averaged the inferred subunit weights for all distant cells to obtain an “average cell” and understand better how this cell type pools stimulation from the far surround (see methods). The subunit weights inside the receptive field center, which implement the linear computation, did not change in the combined bar condition relative to the single distant bar condition (fig. 3C and supp. fig. 1 for OFF subunits). In contrast, the subunit weights pooling the output of distant ON subunits were strongly decreased in the combined bar condition relative to single distant bar condition (fig. 3D). Consequently, stimulation in the receptive field center suppresses the contributions of distant subunits which implement the nonlinear computation. In summary, we observed a switch between two very different computations: cells changed from performing a non-linear computation on distant stimuli to performing a linear computation on the stimuli inside their receptive field centers (fig. 3E).

### Global gain control explains the gradual suppression of distant responses

Our previous results indicated that the influence of distant inputs is suppressed when the receptive field center is stimulated. To elucidate further how central inputs suppress distant ones, we asked if the suppression increases gradually as central inputs become progressively stronger, or if the suppression is only activated once the strength of central inputs exceeds a threshold.

To test this we displayed a series of stimuli where two bars were oscillating over the visual field at incommensurable frequencies (see methods). By averaging over the oscillation period of each bar, we could isolate the responses due to each bar. Our analysis focused on neurons for which one of the bars was within the receptive field center, while the other bar was outside. The central bar was displayed at several luminances, ranging from zero contrast (i.e., at background gray level) to maximally dark bar. We observed that responses to the distant bar decreased gradually as the luminance of the central bar went from gray to full dark, implying that the suppression of distant inputs was gradual (fig. 4A and B). Next, we looked for a general model that could explain center-strength-dependent suppression of responses to distal stimulation.

**Figure 4:**
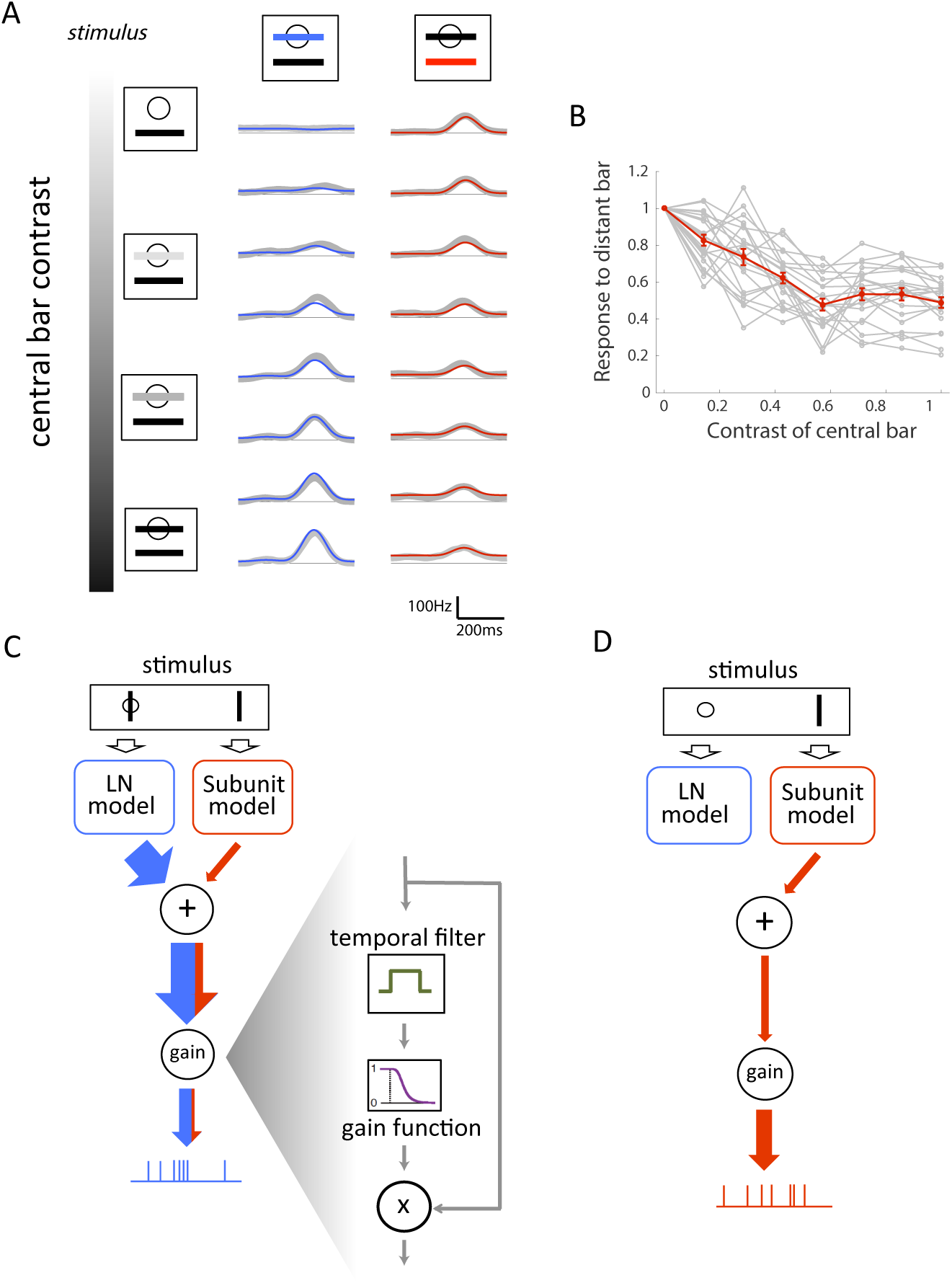
A global gain control model explains gradual suppression of distant responses. **A:** Gradual suppression of the response to the distant bar as central bar contrast increases. Left: schematic of the stimulus. The central bar contrast increased progressively top to bottom. Middle: Response of one cell to the central bar for eight different central bar contrasts; real PSTH (gray) and prediction of the gain control model (blue). Right: Response of the same cell to the distant bar as contrast of the central bar increases; real PSTH (gray) and prediction of the gain control model (red). **B:** Peak of the response to the distant bar (normalized by the response to the distant bar alone) as a function of the contrast of the central bar for individual cells (gray) and on average (red). **C, D:** Schematic of the effect of gain control. In case of a central stimulation (C), a gain control component will suppress the output; weak inputs from the surround will barely modulate the response. In case of no central stimulation (D), the gain control component will enhance the weak inputs corresponding to the distant bar.

We hypothesized that the observed suppression is due to a gain control acting on the ganglion cell. In this view, distant inputs originating in the far surround are much weaker than the inputs originating in the center, and a gain control mechanism normalizes the cell’s firing rate by the total overall input. Specifically, our model sums the inputs coming from central and distant stimulation, averages the result over a long (1 s) temporal window to get the normalization signal, and finally divides the instantaneous input by this normalization to get the final firing rate prediction (see methods). When the center is stimulated, the gain control will thus divide the output by a large normalization factor, which will suppress weak inputs from the surround (fig. 4C). However, when the surround is stimulated alone, the gain control will act as an amplifier, allowing the cell to respond to the distant bar (see fig. 4D for an illustration).

We fitted such a gain control model, inspired by (Shapley and Victor, 1979; Berry et al, 1999), to neurons stimulated by two bars with different luminances (see methods). The model predicted very well the responses to the different stimuli (fig. 4A): it explained 84% ± 3% (n = 168, see methods) of the variance in responses to the distant bar across all contrast conditions. Note that an additive model, where the responses to isolated distant and central bars are simply summed, could not explain any modulation of the distant response by the central bar contrast (fig. 4B).

The subunit model equipped with gain control thus represents a complete functional model able to quantitatively explain the responses of ganglion cells to randomly moving bars. It not only accounts phenomenologically for the suppression of responses to the distant bar due to central motion, but also predicts how such suppression depends on the luminance of the central bar.

### Center computation is position sensitive, while position-independent distant computation codes for large stimulus changes

Our model showed that fast OFF ganglion cells performed two very different computations on the stimulus: a linear one inside their receptive field center and a non-linear one outside. But what visual feature is extracted by each of these computations? To address this question, we first plotted the distribution of bar positions 100 ms before a spike. For an example central cell (fig. 5A), this distribution was narrow and had a cell-dependent preferred location, indicating the ability of central cells to code for the position of the bar. In contrast, for a distant cell the same distribution remained broad and largely overlapping with prior distribution of bar positions (fig. 5A), suggesting that distant cells were largely insensitive to the exact position of the stimulus.

**Figure 5:**
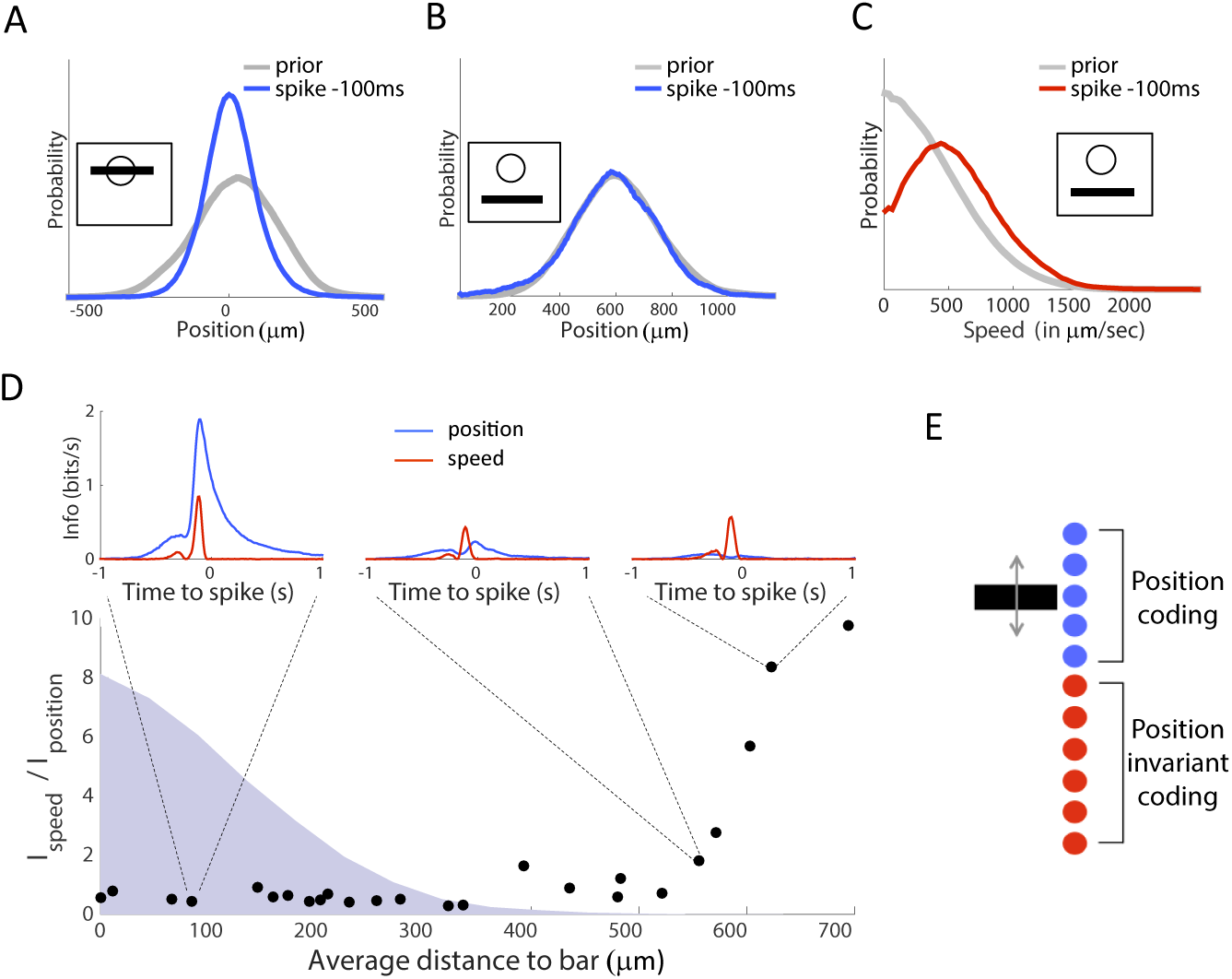
Central computation codes for position, while distant computation is invariant to position and codes for stimulus change. **A:** Distribution of the bar positions for the complete stimulus trajectory (“prior distribution”, gray) and 100 ms before the spike of a central cell (blue). Zero corresponds to the location of the cell’s receptive field (RF) center. **B:** Same as A for a distant cell with its receptive field center far from the bar. **C:** Distribution of the absolute speed of the bar for the complete stimulus trajectory (gray) and 100 ms before the spike of a distant cell (red). **D:** Ratio between the information individual cells carry about bar speed vs about bar position, as a function of the average distance to the bar. Distribution of bar positions is shown as a blue shade. For selected cells, the insets indicate the mutual information between the spiking response and the position (blue) or the speed (red) at different time delays. **E:** Schematic showing that central cells code for bar position while distant cells are nearly invariant to it.

Distant cells nevertheless were selective for the stimulus. The average speed (absolute velocity) preceding the spike of a distant cell showed a preference for fast bar motions (fig. 5C). By quantifying the information carried by each cell about bar position and speed (see methods, n = 25 cells), we confirmed that distant cell responses encoded substantially more information about speed than about position, whereas central cells coded primarily for position (fig. 5D). The observation of highly synchronous responses of distant cells to a random repeated bar trajectory (fig 1) further supports our interpretation that the distant computation is largely invariant to the exact bar position. We also analyzed the subunit model fitted to cell responses in the previous section and found that cells close the bar were much more sensitive to the bar position than distant cells, confirming these results (supp. fig. 2).

Cells close to the bar thus computed a feature strongly related to the exact position of the bar, while distant cells were largely invariant to the bar position (fig. 5E). How can we interpret this invariance to position and simultaneous sensitivity for high-speed motion? Fig 3 shows that cells pooled the output of distant subunits over a large spatial region of the surround. This pooling was largely unselective for position and thus explained how the observed distant responses could remain nearly invariant to the position of the bar. To trigger a response in distant cells, the bar had to sweep across a large region of space: this would lead to an activation of a large number of subunits in a short amount of time, that are summed together to result in the activation of a ganglion cell. This activation should only occur when the bar moves at sufficiently high speed, explaining the preference for high speed motion.

However, our model further predicted that flashing a large object in the cell’s surround should also activate many subunits at the same time and trigger a response. We confirmed this was indeed the case (fig. 6B; note that distant responses are ON responses, consistent with our model, where surround is dominated by ON subunits). This result suggests that distant cells should not be viewed narrowly as encoding “high speed” (an interpretation that is natural for the moving bar stimulus); rather, a generic interpretation is that distant cells code for any “large change” in the stimulus. In summary, central cells are position sensitive, while distant cells are largely insensitive to the exact position of the stimulus and behave like generic “change detectors”.

**Figure 6:**
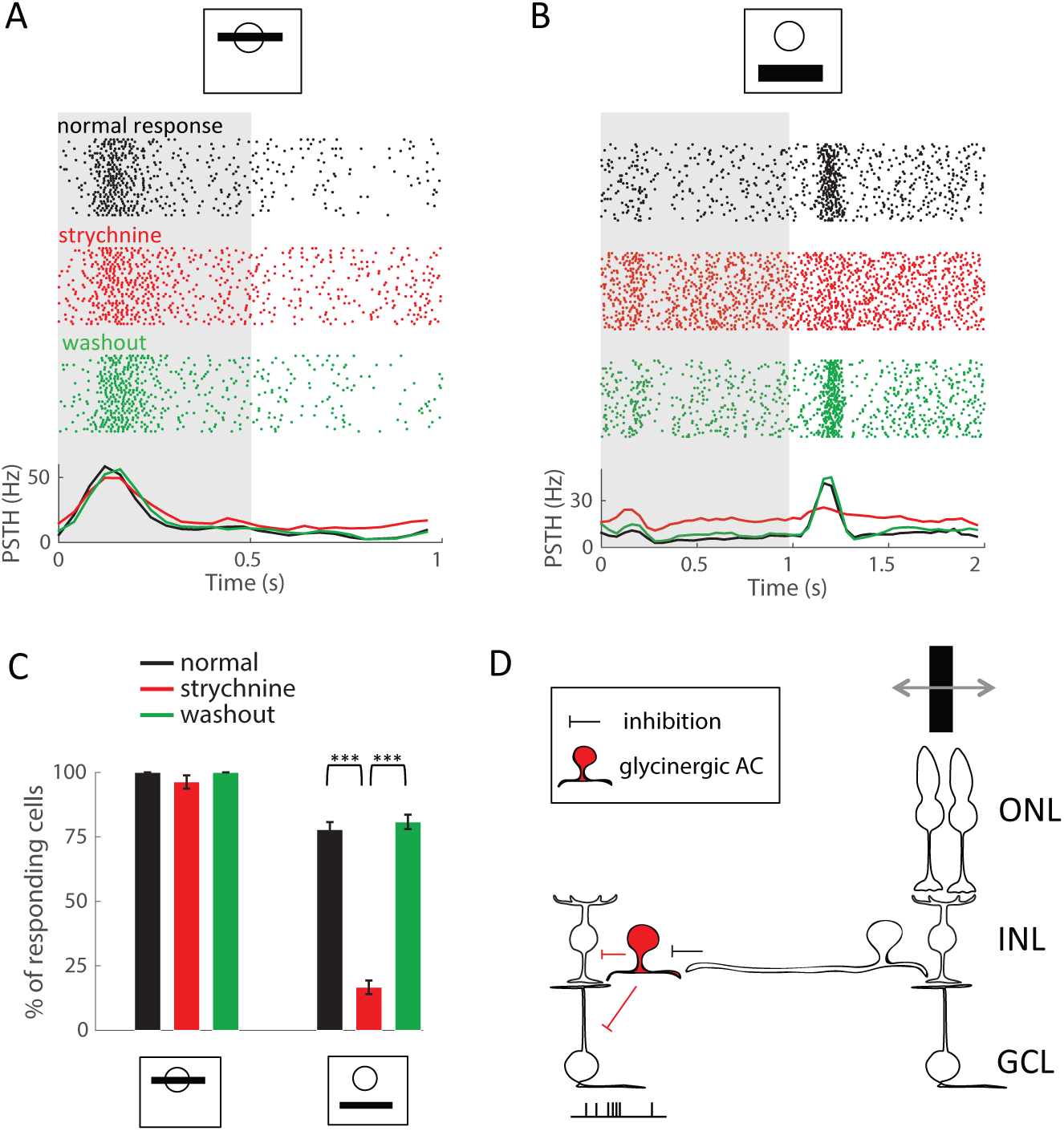
A disinhibitory circuit of amacrine cells is responsible for distant activation. **A:** Response to a bar flashed inside the receptive field center is not altered by the addition of strychnine. Singlecell responses to repeated presentations of the dark bar (shaded gray indicates the time window where the bar was presented); each dot is a spike, each row corresponds to a stimulus repeat. Black raster: control. Red raster: after addition of 1 *µ*M strychnine to the bath. Green raster: after washing out strychnine. Bottom: PSTHs computed from the rasters with the same color code. **B:** Same as A for a bar flashed far from the receptive field center. **C:** Percentage of cells responding to a central flash (left) and to a distant flash (right) before and after strychnine application, and after the drug wash-out. Same colors as **A** and **B**. **D:** Schematic showing a putative circuit for distant activation. When a distant stimulus is present, excitation of amacrine cells inhibits subsequent glycinergic amacrine cells (red), which, in turn, dishinibit bipolar cells and ganglion cells. ONL: outer nuclear layer. INL: inner nuclear layer. GCL: ganglion cell layer.

### A disinhibitory circuit of amacrine cells relays distant inputs

We next examined how the computations required by our phenomenological model could be implemented by the retinal network. The subunits of our model most likely correspond to bipolar cells (Demb et al, 2001; Baccus et al, 2008; Gollisch and Meister, 2010). For subunits in close physical proximity to the ganglion cell, the weights can result from direct synaptic connections between bipolar cells and the ganglion cell. In addition to these proximal connections, however, our model suggested that the ganglion cell also integrated the outputs of distant subunits, albeit with a smaller weight. What could be the circuit basis of such distal integration?

One possible mechanism explaining the activation of ganglion cells by distant stimuli would involve amacrine cells: they could propagate the activity of bipolar cells laterally to distant ganglion cells (Geffen et al, 2007). To test if glycinergic amacrine cells are involved in the distant activation of ganglion cells, we blocked their synaptic transmission with strychnine (see methods). This blocker suppressed distant responses to a flashed bar (fig. 6B, C), while leaving central responses mostly unaffected (fig. 6A, C). Washing out the drug restored the distant responses (fig. 6B, C). Glycinergic amacrine cells therefore constitute a necessary component of the observed distant responses.

Suppression of distant responses by strychnine showed that the weights assigned to distant subunits in our model are mediated by glycinergic amacrine cells. How could these weights be positive, while glycinergic amacrine cells have an inhibitory effect on their post-synaptic targets? One explanation is a disinhibitory loop, where one amacrine cell inhibits its post-synaptic amacrine cell target, which in turn disinhibits the ganglion cell (directly or through bipolar cell) (see also (Manu and Baccus, 2012)). Such a disinhibitory circuit could involve serial connections between GABAergic and glycinergic amacrine cells (Eggers and Lukasiewicz, 2010), or, alternatively, serial connections between different types of glycinergic cells. Glycinergic amacrine cells can ultimately inhibit OFF bipolar cells (Eggers and Lukasiewicz, 2011) or ganglion cells (O’Brien et al, 2003). The net effect of such a disinhibitory circuit is a distant excitation of ganglion cells (fig. 6D).

## Discussion

We have shown that two representations of a stimulus coexist, at the same time, within a neural population formed by ganglion cells of a single type. We constructed a mathematical model that recapitulated the multiplexing of the two relevant computations. To that end, the model required nonlinear summation within the receptive field as well as a gain control mechanism. The model predicted precisely the responses of the fast OFF ganglion cells to a bank of dynamical stimuli which included complex, spatio-temporal stimulation in the far surround. Finally, our experiments suggested that a disinhibitory retinal circuit composed of two amacrine cells could mediate the distant computation.

When an object is moving randomly, neurons whose receptive field centers overlap with the object code for its position, while distant neurons code for general, large-scale changes in the stimulus. Each neuron can switch from one computation to the other depending on the visual context. Recent works have shown that the feature extracted by a cell can change when the average luminance changes (Smirnakis et al, 1997; Tikidji-Hamburyan et al, 2015), or during saccadic exploration of the visual scene (Geffen et al, 2007). Here we show that, in a single visual scene, the same cell type can be used to extract two features *simultaneously*. Feature extraction does not change only with the average luminance of the visual scene. Rather, two features can be extracted at the same time by a single cell type in a single visual scene. These findings expand the traditional view of a “neural map” where there is a one-to-one correspondence between one cell type and one visual feature: here we show that a “neural map” can contain more than one “feature map” at the same time. Multiplexing two computations in a single neural type could enable optimal use of coding resources: if ganglion cells don’t have an object inside their receptive field center, rather than staying silent, they are put to use to code for a different feature of the stimulus.

**Figure 7:**
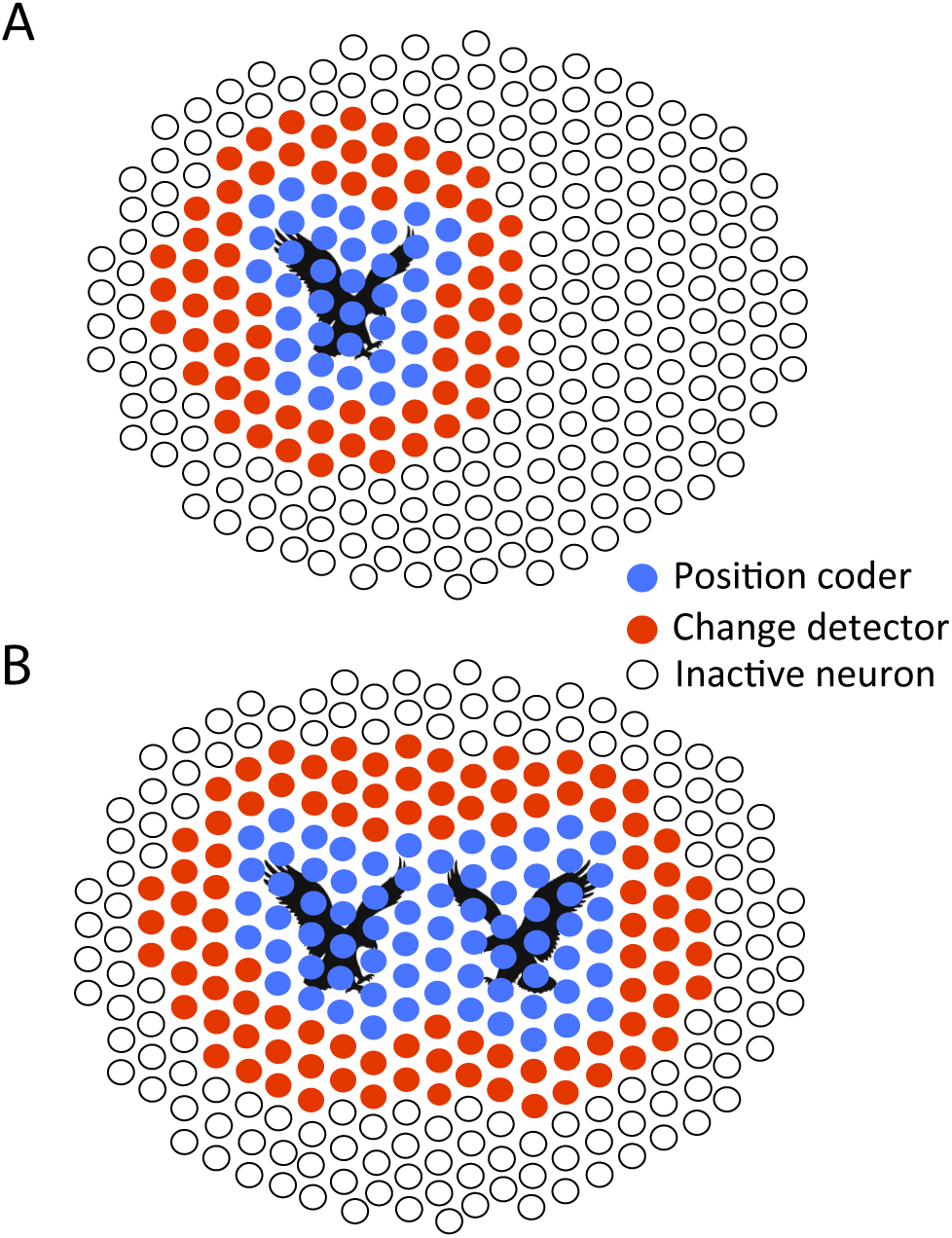
Cells of a single type switch between two different computations. **A:** Schematic of how cells of the fast OFF type code for a moving stimulus (here an eagle depicted in black). Each circle represents a fast OFF cell. Empty circles correspond to inactive cells, red circles to cells acting as change detectors, while blue cells are sensitive to the exact position of the stimulus. **B:** Reorganization in the population code following the simultaneous display of two stimuli. Same legend as A.

Having coexistent stimulus representations present in the same neural map may appear problematic for subsequent stages of processing: how can downstream neurons tease apart spikes corresponding to one or the other computation? Our study shows that a large ensemble of ganglion cells will respond synchronously and sparsely to distant objects. While such responses could appear negligible or ambiguous when observed at the single cell level, access to the complete population would enable downstream processing to unambiguously recognize distant responses by detecting synchronous ganglion cell activity. Thus, synchrony could serve as a signature in the spike trains to separate position signals from change signals.

Previous works have shown that ganglion cells can be activated by fast motion in their far surround (“shift-effect”: (McIlwain, 1964; Cleland et al, 1971; Ikeda and Wright, 1972; Fischer et al, 1975; Barlow et al, 1977)). Here we constructed a model that can accurately predict how fast OFF ganglion cells would respond to distant, complex stimuli, and how these distant stimuli would be integrated with other stimuli simultaneously displayed inside the receptive field center. Previous models mostly focused on how the surround modulates responses to central stimuli. However, how responses to distant stimuli can modulate ganglion cells themselves, and how they could be affected by center stimulation, has received less attention (Shapley and Victor, 1979). Demb et al (1999) found that inputs from center and surround stimulation were summed linearly, while we found a non-linear suppression of distant inputs. This discrepancy could be due to a difference of species, cell type, or recording technique. Passaglia et al (2001) showed that distant stimulation could be suppressed by center stimulation, but the timescale of the modulation was much longer than in our work. Interestingly, Jadzinsky and Baccus (2015) suggested a model to predict how stimulation of the surround can affect the selectivity to the center stimulation that bears some similarity with our model. In most studies, the stimulus employed to modulate activity from the surround was very large. In our study, we showed that the *same* stimulus triggered two different types of responses, a central one and a distant one, within the same type of ganglion cell, demonstrating the coexistence of the two representations.

Our results suggest that the retinal network implements the activation of ganglion cells by distant stimuli through a disinhibitory circuit in which intermediary amacrine cells are activated by bipolar cells and subsequently inhibit glycinergic amacrine cells. This release of glycinergic inhibition can affect both OFF bipolar cells (Eggers and Lukasiewicz, 2010) and OFF ganglion cells (O’Brien et al, 2003), and results in OFF ganglion cell activation. It is unclear if this disinhibitory relay is composed of GABAergic and glycinergic cells, or only of glycinergic cells. Attempts to disentangle the two hypotheses by blocking GABAergic transmission triggered large oscillations in the retina, making the results difficult to interpret (Demb et al, 1999). A similar disinhibitory circuit might also be involved in other kinds of complex processing taking place in the ganglion cell surround. When large visual features stimulate distant regions of the surround, the inhibitory input to bipolar cells (Eggers and Lukasiewicz, 2010) and ganglion cells (O’Brien et al, 2003) was reduced. This reduction of surround inhibition was mediated by a disinhibitory circuit similar to the one we uncovered.

We have shown that a single cell type mosaic can simultaneously multiplex several fundamentally distinct computations. Our findings considerably enrich the classical view of ganglion cell types as being tightly linked to their corresponding feature maps, and uncover the flexibility of the retinal code when stimulated with complex, dynamical stimuli. The notion of a feature map is central to most sensory structures. Flexible computations, where several features are represented by a cell type simultaneously in response to complex stimuli, might also be implemented in other sensory areas. It remains to be understood whether this flexibility can be seen as arising from some efficient coding principle (Tkaˇcik and Bialek, 2016), and how such flexible coding schemes can be interpreted by the downstream areas (Botella-Soler et al, 2016).

## Material and methods

Unless stated otherwise, all error bars in figures and text are standard error of the mean (SEM). SD stands for standard deviation.

### Retinal recordings

Recordings were performed on the Long-Evans adult rat. Animals were euthanized according to institutional animal care standards. The retina was isolated from the eye under dim illumination and transferred as quickly as possible into oxygenated AMES medium. The retina was then lowered with the ganglion cell side against a multi-electrode array whose electrodes were spaced by 60 *µ*m, as previously described (Marre et al, 2012; Yger et al, 2016). Raw voltage traces were digitized and stored for off-line analysis using a 252-channel preamplifier (MultiChannel Systems, Germany). The recordings were sorted using custom spike sorting software developed specifically for these arrays (Marre et al, 2012; Yger et al, 2016). We extracted the activity of a total of 810 neurons over 5 experiments with satisfying standard tests of stability and limited number of refractory period violations.

### Visual stimulation

Our stimulus was composed of one or two black bars moving randomly on a gray background. Each bar was animated by a Brownian motion, with additional feedback force to stay above the array, and repulsive forces so that they do not overlap. The bars stayed within an area that covers the whole recording array. The amplitude of the bar trajectories allowed them to sweep the whole recording zone. The trajectories of the bars x_1_ and x_2_ are described by the following equations (Mora et al, 2015):

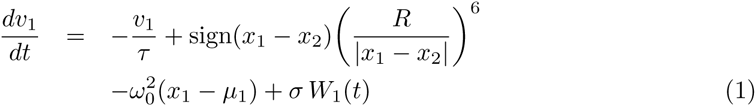

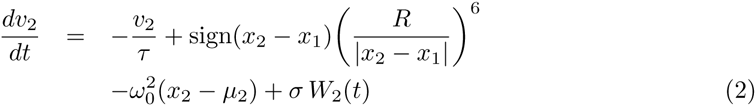

where *W*_1_(t) and *W*_2_(t) are two Gaussian white noises of unit amplitude, *µ*_2_ *µ*_1_ = 600*µ*m is the shift between the means, *ω*_0_ = 1.04 Hz, *τ* = 16.7 ms, R = 655*µ*m and σ = 21.2*µ*m • s^−3/2^. The width of one bar is 100*µ*m. The stimulus was displayed using a Digital Mirror Device and focused on the photoreceptor plane using standard optics. For receptive field mapping, a random binary checkerboard was displayed for 1 hour at 50 Hz (check size: 60 *µ*m).

All the other stimuli used (for classification of cells, fitting the gain control model and pharmacological study) are described in the corresponding method section. For all stimuli, the level of light of the gray background was between 10^12^ and 10^13^ photons.cm^−2^.s^−1^.

### Typing

We performed cell classification based on the response of the cells to a set of stimuli and on their temporal receptive field.

*Full field flicker*: this stimulus consisted of a 15-seconds sequence of a full-field stimulus, repeated 100 times. The stimulus was generated by selecting a random row of pixels from a natural image and displaying subsequently at 40Hz the intensity of these pixels uniformly on the entire screen.

*Shifting barcode*: this stimulus consisted of an alternation of white and black stripes of width 70 *µ*m chosen randomly, moving at a constant speed of 1000 *µ*m/s in the 4 cardinal directions. For each direction, the 17-seconds sequence was repeated 30 times.

For each cell, we created a vector by concatenating the PSTH in response to the *full field flicker* stimulus, the 4 PSTHs in response to the *shifting barcode* stimulus corresponding to the 4 cardinal directions, the temporal receptive field and the auto-correlogram of the cell in response to the checkerboard stimulus. The PSTHs of the *shifting barcode* were temporally realigned beforehand according to the receptive field location of each cell. PSTH for each stimulus was normalized such that they all had a mean of 0 and a variance of 1.

We then performed PCA on this collection of vectors. We kept the projections on the first eigenvectors in order to explain 95% of the total variance. We then performed clustering on these vectors using the peak density algorithm (Rodriguez and Laio, 2014). The threshold parameters of the algorithm were manually adjusted in order to select the outliers as centroids of the clusters. This method allowed us to identify reliably an OFF type of ganglion cells across all experiments. The receptive fields (RF) were regularly tiling the visual field, with little overlap between them. This mosaic property, often observed in the retina, was used here as a validation of our typing procedure, as we did not use the position of the RFs in the clustering procedure.

### Synchrony between cells

To quantify the synchrony between cells, we displayed a 10-second bar movie to the retina, repeated 54 times. A maximum of 25 cells of the same type recorded simultaneously were subdivided in two groups, the distant cells, that were more than 200 *µ*m away from the central bar position, and the central cells, that were less than 200 *µ*m away from the central bar position. For all cells we computed the PSTH with a time bin of 20 ms. We computed the Pearson coefficient between all pairs of PSTHs of distant cells, and all pairs of PSTHs of central cells respectively. We grouped the pairs based on the distance between their receptive field centres along the bar motion axis.

### Linear model and subunit model

#### Subunit model

The subunit model is a two-layer model that predicts the response of a ganglion cell to the moving bar. Each layer performs a linear combination of its inputs followed by a non-linear transformation. The first layer is a collection of identical and translated Linear-Non-Linear (LN) units. The second layer is a unique LN unit taking the output of the first layer as an input.

In the first layer, we tiled the space with 200 bipolar-like ON and OFF subunits on a onedimensional lattice, with subunits equally spaced at 20*µ*m interval. Each unit had a receptive field with a Gaussian spatial profile of the right polarity and a biphasic temporal profile, modelled by a sinusoid. All units of a same polarity are identical up to a translation. The non-linearity was a rectified square function, h. The output of the first layer was therefore:

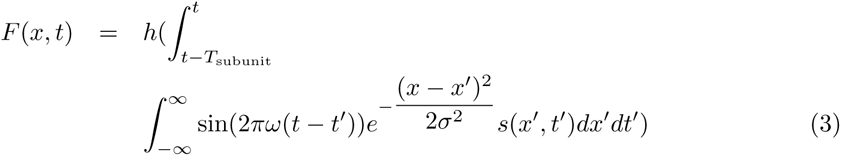

where *h*(*x*) = *x*^2^ if *x* ≥ 0, and 0 otherwise. *T*_subunit_ = 0.3 s, *ω* = 1/*T*_subunit_, σ = 30 μm

The stimulus movie s(x, t) was one-dimensional in space because the stimulus was a long bar, whose length can be considered infinite. We used a temporal binning of 17ms, corresponding to the refresh rate (60Hz) of the screen used to project the movie on the retina.

The second layer consisted of a single Linear-Non-Linear Poisson unit. The unit pooled linearly its inputs from all the subunits of the first layer according to a kernel K, with an extension in time of 0.5 seconds. To obtain the firing rate r(t) of the cell, the weighted sum was passed through a non-linearity of the form f(x) = log(1 + exp(x)). The spikes were then generated according to a Poisson process.

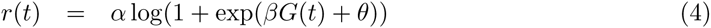

where

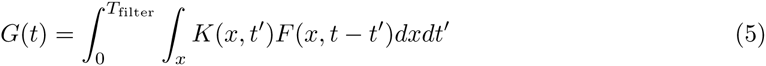

with *T*_filter_ = 0.5 s, and *α, β, θ* are parameters of the non-linearity that are fitted to the data. The linear model (LN) was built using the same architecture as the subunit model, except that the rectified square non-linearities in the subunits were replaced by the identity.

### Fitting

For both models we used the same fitting procedure. The parameters of the kernel K and the parameters of the spiking non-linearity *α, β, θ* were the only parameters fitted to the data. The kernel parameters and the spiking non-linearity parameters were fitted alternatively using block gradient descent (McFarland et al, 2013) across 6 iterations. The repeated parts of the stimulus were held back during fitting and were used to cross-validate the model.

The parameters of the kernel were optimized to maximize the log-likelihood function of the spike train under Poisson assumption (McFarland et al, 2013). For this optimization we performed Limited-memory BFGS gradient descent on the parameters of the kernel (McFarland et al, 2013). In order to avoid overfitting, we imposed two regularisation constraints: spatiotemporal smoothness and sparseness of the kernel. The cost function C was of the form:

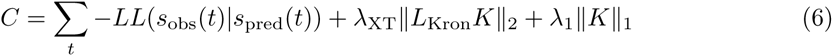

where *LL* is the loglikelihood of the observed spike train *s*_obs_ (under Poisson assumption), *K* is the kernel defined above, *λ*_XT_ = 300 is the penalty term enforcing smoothness of the kernel, L_Kron_ is the Kronecker sum of discrete Laplacians, *λ*_1_ = 400 is the L1 penalty term enforcing sparseness of the filter coefficients.

The penalty terms were chosen to minimize overfitting. To fit the linear model (LN), we divided by 10 these two penalty terms as it slightly improved the performance of the model for distant cells. The parameters of the non-linearity were fitted by minimizing the cost function with the active-set method. The following constraints were enforced: *α >* 0, *β >* 0, *θ* has an upper bound. *β* and θ were redundant with the kernel parameters but adding them accelerated the convergence of the optimization (McFarland et al, 2013).

### Quantification of the performance of the LN model and of the subunit model

We fitted the model on the unrepeated part of the stimulus and we tested the performance of the model on the repeated part of the stimulus (54 repetitions of a 10 second sequence). For each cell we then computed the Pearson coefficient r between the real PSTH and the predicted PSTH (time bin: 17 ms). Population averages are indicated in the text as mean ± standard error of the mean. In figure 2F, we set to zeros all negative Pearson coefficients for readability. In order to show that the LN model was performing significantly better for central stimulation than for distant stimulation, we selected only the cells that were less than 300 *µ*m away from the bar in one condition and more than 400 *µ*m away from the bar in the other condition. We then performed a paired t-test comparing the performance of the LN model in both conditions for each cell.

A possible explanation for why the linear model performed poorly for distant cells could be that distant stimulation evoked less reliable responses. In order to exclude this possibility, we computed the ratio of explainable variability predicted by the model. The explainable variability was defined as the average Pearson coefficient between pairs of PSTHs generated by instantiations of a Poisson process with mean firing rate equal to the real firing rate of the cell estimated from the PSTH. We divided the performance of our model (defined as the Pearson coefficient between real and predicted PSTH) by this explainable variability to obtain the ratio of explainable variability predicted by our model.

### Calculation of the average linear filters in the subunit model

To compute the average filter in the one-bar condition (fig 3D), we selected only the cells stimulated outside of their receptive field (RF) center. Our criterion was that the bar central position should be more than 200 *µ*m away from the RF center.

To compute the average filter in the two-bar condition (fig 3C), we selected only the cells that were stimulated inside their receptive field centers by at least one of the bars. Our criterion was that the bar central position should be less than 200 *µ*m away from the RF center.

For all cells and in both bar conditions, only a portion of the extended receptive field center was visited by the bars, therefore inducing a bias in the filters fitted on these movies. To compute the weights of the average filter without bias, we first realigned the filter of each cell relative to the center of its receptive field. Then for each coordinate (x,t) of the average filter we averaged the corresponding subunit weights for the subset of cells for which the coordinate was visited more 200 times/hour by the bar.

### Suppression index

In figure 3B, we quantified the suppression of the response to the distant bar when there was another bar moving inside its receptive field center. For this we defined the residual response to the distant bar in case of a central bar as:

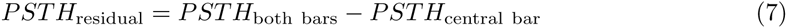

We then computed the suppression index, defined as:

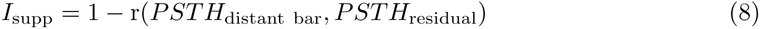

where *r* is the Pearson coefficient. If the suppression of the distant bar response is complete, the index should be equal to one. If there is no suppression, and the responses to each bar are summed, then the index should be equal to 0 in the absence of noise. However, since noise is present, we defined a suppression index for the linear model, which reflects the index value that should be expected purely from noise, without suppression of the distant response:>

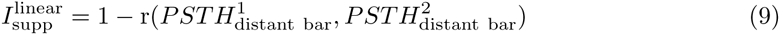

where *PSTH*^1^_distant bar_ and *PSTH*^2^_distant bar_ were computed on two different sets of trials. We performed this quantification on the 25 cells recorded and plotted the mean and SEM of the suppression index for the real data and for the equivalent linear model. A suppression index higher than 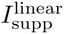 indicates a true suppression that cannot be explained by noise.

### Gain control model

We displayed two bars of width 300 *µ*m and separated by 800 *µ*m, oscillating with a sine wave trajectory at slightly different frequencies: the central bar was oscillating at 2 Hz and the distant bar at 1.98 Hz. The central bar was played at 8 different contrasts interleaved randomly and ranging linearly from 0 to 1. For each contrast, the two bars were oscillating during an uninterrupted sequence of 50 seconds, so that the central bar had traveled exactly 100 periods and the distant bar exactly 99 periods during a sequence. At the end of a sequence, all possible phase shifts between the two bars had been visited exactly once. This trick allowed us to average out the influence of one bar when computing the PSTH on the period of the other bar.

To show the gradual suppression of the distant response in fig. 4B, we normalized the amplitude of the response to the distant bar by the amplitude of the response to the distant bar alone (i.e. zero contrast for the central bar).

We then fitted a single model on all contrast conditions. The model was of the form:

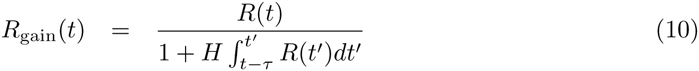

where *τ* = 1 s is the time constant of integration of the gain control and H is the gain. R(t) is the total response before application of the gain control, given by the equation:

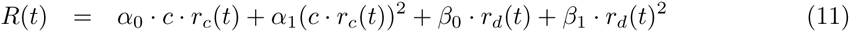

where *r_c_* is the response to the central bar alone at full contrast, *r_d_* the response to the distant bar alone, c is the contrast. *r_c_* and *r_d_* were estimated from the PSTHs in response to the central bar and to the distant bar played alone respectively. We needed to introduce quadratic terms because the PSTH for the central bar condition depended quadratically on the contrast of the central bar. This is consistent with our subunit model, where the first layer contained a rectified quadratic function h.

We fitted the parameters *α*_0_, *α*_1_, *β*_0_ and *β*_1_ and *H* so as to maximise the log-likelihood of the spike train under Poisson assumption (bin size: 17 ms). To adjust the parameters we used the active set method. However, we fixed the parameter *τ* to 1 second because the periodicity of the stimulus did not allow us to explore thoroughly the time constant of integration of the gain. To test our model, we measured for each cell (n=21) and each contrast the amplitude of the response to the distant bar (defined as max(PSTH)-min(PSTH), bin: 100 ms) and compared it to the amplitude predicted by our model. We then estimated the percentage of variance explained by our model across all cells and conditions using bootstrapping.

### Information estimation

The information conveyed by the cell response R about the stimulus X (i.e. mutual information between R and X) is equal to the reduction in entropy of the distribution of X provided by the knowledge of R.

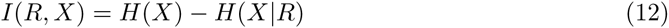

In our case we first defined the stimulus as the position *P (t + δt)* of the moving bar for different lags δt relative to the cell response *R*(*t*) (in fig. 4C, δt is the x-axis of the insets). The lags were introduced to account for the delay in the neural response. We discretized linearly the space of P in 10 bins in order to have a well-sampled distribution with our finite dataset. We discretized the spike train in 10 ms bins and we binarized it by setting to 1 all the bins where there was at least one spike and to 0 the other bins. Changing the discretization steps used to bin P and the spike train did not change qualitatively our results. Then we computed the mutual information between the cell response and the instantaneous position of the bar with a lag δt ranging from -1 second (information about the past stimulus) to 1 second (information about the future stimulus):

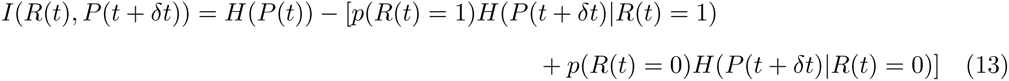

Note that the information about the future of the stimulus was not always zero. This is because the successive positions of the bar are correlated in time, so that part of the information conveyed by the cell response about the past position of the bar is also informative about the future position of the bar. We then defined the stimulus as the speed of the bar S with different lags δt relative to the cell response. The speed was defined as:

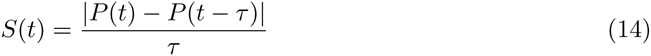

where *τ* = 100 ms. We discretized linearly the space of S in 10 bins and we computed mutual information between *R*(*t*) and S(t + δt). To estimate the information rate in the insets of figure 5D, we divided the mutual information by the bin size (10 ms). For each cell, we finally computed the ratio between the maximum of *I(*R*(*t*), S(t + δt*)) and the maximum of *I(*R*(*t*),P (t + δt*)) over all time lags tested.

### Pharmacology

To block glycinergic transmission, we added 1 microMol strychnine (Sigma-Aldrich ref. S8753) to the bath (Curtis et al, 1971; Schaeffer and Anderson, 1981; Lee et al, 2016; Menger and Wassle, 2000). To generate the rasters and PSTHs in response to the central bar, we flashed a dark bar of width 100 *µ*m in the center of the receptive field of the cell for 0.5 s 40 times, separated by 0.5 s of gray screen. For the distant responses, we used 230 *µ*m wide bars flashed for 1 s, in a region 0.5 to 1 mm away of the cell’s receptive field center. For the population analysis, we flashed a bar 100 *µ*m wide in random locations relative to the receptive fields of the cells, 20 times at each location. For each cell recorded of the type under study (17 cells), we selected the flashes that were less than 80 *µ*m away from the receptive field center to study the effect of central stimulation. To study the effect of distant stimulation, we selected the flashes that were between 200 *µ*m and 500 *µ*m away from the cell receptive field center. For each stimulus and each cell, significant responses were determined based on a z-score analysis. We estimated the mean and standard deviation (SD) of the activity prior to stimulus and considered that a response was detected if the activity exceeded the mean by more than five times the SD in the second following the onset of the stimulus (for a bin size of 40 ms). To estimate the percentage of responding cells in fig. 6, we estimated means and standard errors of mean by pooling together all stimulus conditions across all the cells. We performed a one-tailed two-sample t-test to assess the reduction of responses to the distant flash after drug was added to the bath. The p-value was less than 10^*−*3^.

## Acknowledgments

We thank Vicente Botella-Soller, David Kastner, Stuart Trenholm and Michael J. Berry II for fruitful discussions, Valerie Fradot for excellent technical support, Hanen Kabou for her illustrations, and Thierry Mora for critical reading of the manuscript. This work was supported by ANR-14-CE13-0003 to P.Y., ANR TRAJECTORY, ANR OPTIMA, the French State program Investissements d’Avenir managed by the Agence Nationale de la Recherche [LIFESENSES: ANR-10-LABX-65], a EC grant from the Human Brain Project (FP7-604102)), NIH grant U01NS090501 to OM, a Foundation Figthing Blindness grant to SP, and the Austrian Research Foundation FWF P25651 to GT. S.D. was supported by a PhD fellowship “DIM cerveau et pensee” from the region Ile-de-France.

## Supplementary Figure 1: OFF subunit weights

**Supplementary Figure 1:**
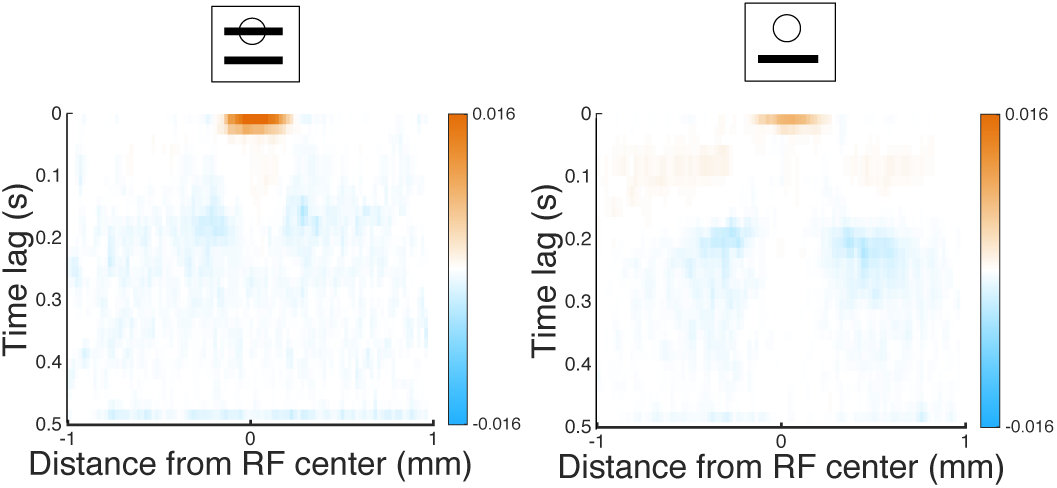
**A, B:** Spatio-temporal distribution of the OFF-subunit weights for the second stage of the subunit model, averaged over all cells. **A:** combined bar stimulus. **B:** a single distant bar stimulus.

## Supplementary Figure 2: Sensitivity of the model to changes in the absolute position

Our results show that cells close to the bar were much more sensitive to the bar position than distant cells. Here we show that the subunit model fitted on the cell responses also had this property. For this we directly used our model to estimate the amount of information about a change in the absolute position of the bar trajectory.

To determine the sensitivity of each cell to a change in the absolute position we estimated the KullbackLeibler divergence d_KL_(∆x)between the cell response to an initial trajectory *x*(*t*), and the response to the same trajectory displaced by a small constant shift ∆x. We picked randomly a time T in the stimulus trajectory, and extracted the trajectory *x*(*t*) of the bar for t between T and T + DT (in the following DT = 16 s but the exact value did not change significantly the results). For each cell we then estimated d_KL_(∆x) between the model response to *x*(*t*) and the model response to *x*(*t*)+∆*x*(*t*), where ∆*x*(*t*) = constant is a uniform perturbation of the trajectory, for *t ϵ* [*T,T*+*DT*]. We repeated this estimation many times for different times T (each point in the scatter plots of supp. fig. 2 corresponds to one cell and one choice of T).

To estimate d_KL_(∆x) we assumed that ∆x is small, so that we can expand the Kullback-Leibler divergence up to the second order to obtain:

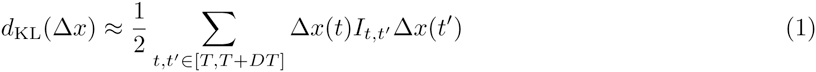

where the matrix I_t,t_*0* is the Fisher Information Matrix of the response distribution conditioned to the stimulus:

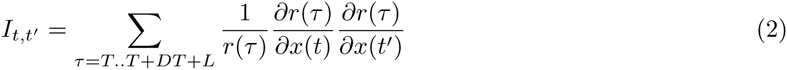

where *x*(*t*) is the position at time t and r(*τ*) is the firing rate at time *τ* predicted by the subunit model in response to the stimulus. L = 0.5 s corresponds to the maximal latency of the response to the stimulus. We then defined the sensitivity as d_KL_(Δ*x*) for a normalized perturbation such that Ʃ_t_∆*x*(*t*)^2^ = 1. We estimated this quantity for all the cells where the model had a very good prediction performance (r ≥ 0.7 in fig. 2F).

**Supplementary Figure 2:**
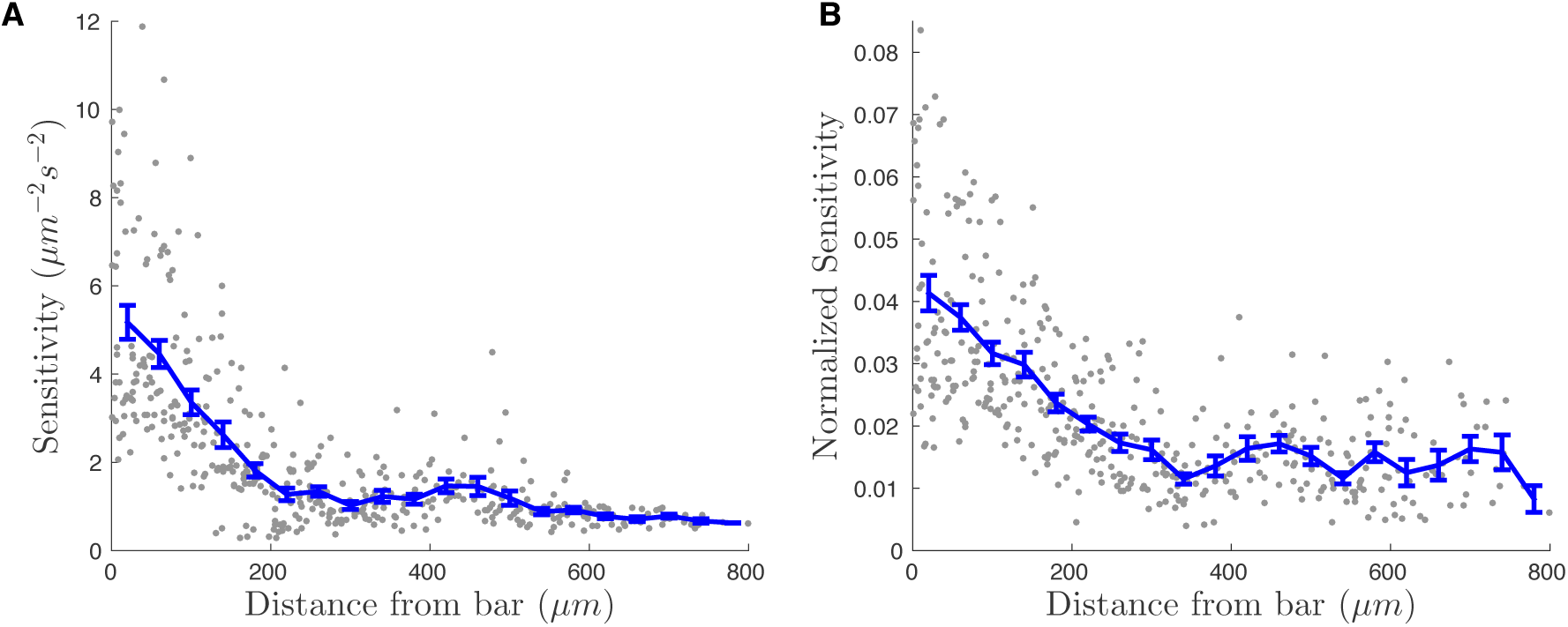
**A:** Sensitivity (see text for definition) of cells to a change in the absolute position of the stimulus as a function of the distance of the cell to the bar. Each point corresponds to one cell and one choice of T (see text). **B:** Normalized sensitivity (see text for definition) of cells to a change in the absolute position of the stimulus as a function of the distance of the cell to the bar.

For cells close to the bar, sensitivity to changes in the absolute position of the bar was high and strongly decreased for distant cells (supp. fig. 2A). We then asked if this decrease is specific to this uniform perturbation, or if it is a global decrease of sensitivity of distant cells to any perturbation. To test this we estimated the maximal sensitivity of each cell, which is the largest eigenvalue of the Fisher information matrix I_t,t_*0*. We normalized the previous sensitivity values by this maximal sensitivity to obtain a “normalized sensitivity”. Even after this, we observed a decrease of this normalized sensitivity with distance (supp. fig. 2B). These results show that the model fitted on the cells has the same property than found on the data previously: central cells were much more sensitive to stimulus position than distant cells.

